# Spatial patterning of tissue volume loss in schizophrenia reflects brain network architecture

**DOI:** 10.1101/626168

**Authors:** Golia Shafiei, Ross D. Markello, Carolina Makowski, Alexandra Talpalaru, Matthias Kirschner, Gabriel A. Devenyi, Elisa Guma, Patric Hagmann, Neil R. Cashman, Martin Lepage, M. Mallar Chakravarty, Alain Dagher, Bratislav Mišić

**Affiliations:** McConnell Brain Imaging Centre, Montréal Neurological Institute, McGill University, Montréal, Canada; Integrated Program in Neuroscience, McGill University, Montréal, Canada; Cerebral Imaging Center, Douglas Mental Health University Institute, McGill University, Montréal, Canada; Department of Psychiatry, McGill University, Montréal, Canada; Department of Biological and Biomedical Engineering, McGill University, Montréal, Canada; Department of Psychiatry, Psychotherapy and Psychosomatics, Psychiatric Hospital, University of Zurich, Switzerland; Department of Radiology, Lausanne University Hospital (CHUV-UNIL), 1011 Lausanne, Switzerland; Department of Medicine (Neurology), University of British Columbia, Vancouver, Canada

**Keywords:** connectome, schizophrenia, intrinsic networks, disease epicenter, anterior cingulate, ventral attention network

## Abstract

**Background:** There is growing recognition that connectome architecture shapes cortical and sub-cortical grey matter atrophy across a spectrum of neurological and psychiatric diseases. Whether connectivity contributes to tissue volume loss in schizophrenia in the same manner remains unknown.

**Methods:** Here we relate tissue volume loss in patients with schizophrenia to patterns of structural and functional connectivity. Grey matter deformation was estimated in a sample of *N* = 133 individuals with chronic schizophrenia (48 female, 34.7 ± 12.9 years) and *N* = 113 controls (64 female, 23.5 ± 8.4 years). Deformation-based morphometry (DBM) was used to estimate cortical and subcortical grey matter deformation from T1-weighted MR images. Structural and functional connectivity patterns were derived from an independent sample of *N* = 70 healthy participants using diffusion spectrum imaging and resting-state functional MRI.

**Results:** We find that regional deformation is correlated with the deformation of structurally- and functionally-connected neighbours. Distributed deformation patterns are circumscribed by specific functional systems (the ventral attention network) and cytoarchitectonic classes (limbic class), with an epicenter in the anterior cingulate cortex.

**Conclusions:** Altogether, the present study demonstrates that brain tissue volume loss in schizophrenia is conditioned by structural and functional connectivity, accounting for 25-35% of regional variance in deformation.

## INTRODUCTION

The human brain is a complex network of anatomically connected and perpetually interacting neuronal populations. The connectivity of the network promotes interregional signaling, manifesting in patterns of synchrony and co-activation [5, 29]. The network also supports molecular transport, including molecules and organelles required for neurotransmission and metabolism [50]. Altogether, the architecture of this white matter “connectome” fundamentally shapes the development and function of the brain [28].

Despite many benefits for communication efficiency and resource sharing, networked systems are also vulnerable to damage [26]. Connections among elements allow pathological perturbations to spread between multiple nodes. Multiple neurodegenerative diseases, including Alzheimer’s and Parkinson’s diseases, are increasingly conceptualized as arising from trans-neuronal spread of pathogenic misfolded proteins [36, 74]. As a result, patterns of neurodegeneration resemble the underlying structural and functional architecture, and are often centered on one or more specific epicenters [60, 77, 80].

Widespread reductions in tissue volume are also observed in schizophrenia (hereafter referred to as *deformation*), but their origin remains unknown [38, 63, 65, 67, 68]. Patterns of cortical thinning in schizophrenia are highly organized and circumscribed by specific networks [46, 67], raising the possibility that connectome architecture shapes the pathological process. Anatomical connections may potentially allow pathogens, such as misfolded proteins or inflammatory markers, to propagate between regions [26]. Additionally, transport of trophic factors between regions may be disrupted if white matter projections are compromised. Both mechanisms would lead to cortical and subcortical deformation patterns that reflect the white matter architecture (i.e., white matter connectivity patterns). Several recent studies have found evidence consistent with this idea. For example, cortical thinning and infracortical white matter breakdown (as measured by fractional anisotropy) appear to be concomitant in patients with schizophrenia [23] and in patients with *de novo* psychotic illness [42]. In addition, cortical thinning is more pronounced among regions with stronger grey matter covariance [73]. Altogether, there is considerable - though circumstantial - evidence that connectivity influences deformation patterns.

Here we test the hypothesis that distributed deformation patterns in schizophrenia are conditioned by connectome architecture. We first estimate cortical and subcortical grey matter deformation in a sample of patients with chronic schizophrenia. We also derive structural and functional networks from independent samples of healthy participants. We then investigate whether deformation patterns are more likely to be concentrated within intrinsic networks, and whether regions that experience greater deformation are also more likely to be connected with each other.

## METHODS and MATERIALS

### Discovery dataset: NUSDAST

Data were downloaded from the Northwestern University Schizophrenia Data and Software Tool (NUSDAST) [37], via XNAT Central (http://central.xnat.org/) and the SchizConnect data sharing portal (http://schizconnect.org/). Detailed information about the data collection procedure is available in [37]. Data comprised 1.5T T1-weighted MRI scans for the baseline visit of 133 individuals with schizophrenia (48 female, 34.7 ± 12.9 years) and 113 age- and sex-matched healthy controls (64 female, 23.5 ± 8.4 years).

### Replication dataset: Douglas Institute

Data comprised 3T T1-weighted MRI scans of *N* = 108 individuals with schizophrenia (26 female, 35.2 ± 8.2 years) and *N* = 68 age- and sex-matched healthy controls (21 female, 34.1 ± 9.0 years) in the independent replication dataset. Detailed information about the data collection procedure is available in [6].

### Regional brain deformation

Local changes in brain tissue volume density were calculated using deformation-based morphometry (DBM; [3]). Regional DBM values are estimated from the deformation applied to each voxel to non-linearly register each MRI scan to a standard template and can be used as measures of tissue loss (termed as *deformation* in this manuscript) or tissue expansion [15, 18, 39, 64, 80]. For detailed information about the DBM procedure, please see *Supplemental Information*. We defined schizophrenia-related deformation patterns using a two-tailed two-sample *t*-test of voxel-wise differences between DBM values of patients and controls, while controlling for age, to assess the group effect on deformation maps. *T*-statistics were converted to *z*-statistics and FDR corrected (5% alpha; [7]), where a larger positive *z*-score corresponds to greater deformation in patients.

### Network reconstruction

#### Anatomical atlas

The brain was parcellated into 68 cortical areas according to the Desikan-Killiany atlas [22]. The parcels were then further subdivided into 114, 219, 448 and 1000 approximately equally sized parcels [14] (referred to as “Cammoun atlas” throughout the present report). The deformation value of each parcel was estimated as the mean deformation of all the voxels assigned to that parcel. Analyses were repeated at all five parcellation resolutions to ensure effects are independent of spatial scale.

#### Healthy structural and functional networks

Structural and functional connectivity data, collected from 70 healthy individuals (age 28.8 ± 9.1 years, 27 females) on a 3T scanner and described in detail elsewhere ([33, 45] (http://doi.org/10.5281/zenodo.2872624), were used to construct high-quality reference brain networks. Deterministic streamline tractography was used to construct structural connectivity matrices for each healthy individual from their diffusion spectrum imaging (DSI). A binary group-average structural connectivity matrix was generated using a consensus approach preserving the edge length distribution in individual participants [8, 10, 44, 45]. Eyes-open resting-state functional MRI (rs-fMRI) scans were collected in the same participants and pre-processed as described in [45]. Functional connectivity between pairs of brain regions was estimated as a zero-lag Pearson correlation coefficient between regional rs-fMRI time series. A group-average functional connectivity matrix was estimated as the mean connectivity of pair-wise connections across individuals. For more detail, please see *Supplemental Information*.

#### Neighbourhood deformation estimates

Structural and functional networks were used to define the neighbours of each brain region. The collective deformation of structural neighbours of *i*-th brain region (*D*_*i*_) was estimated as the mean deformation of all the brain regions that are connected to node *i* by a structural connection (i.e., the nodes with no structural connection to the node under consideration were excluded):

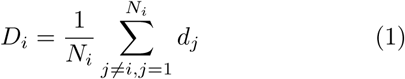

where *D*_*i*_ is the collective deformation of structural neighbours of the *i*-th node, *d* is the node deformation, and *N*_*i*_ is the total number of nodes that are connected to node *i* with a structural connection in the group-level structural connectivity network (i.e., degree of node *i*). The correction term 1/*N*_*i*_ was added to normalize the sum by the number of connections (i.e., correcting for node degree). Note that the summation excludes the deformation value of the node under consideration (*j* ≠ *i*). Altogether, a single value was estimated as the mean neighbour deformation for each node and a correlation coefficient was calculated between the nodes’ and the neighbours’ mean deformation values.

The collective deformation of functional neighbours of node *i* (*D*_*i*_) was estimated in an analogous manner, but the deformation values of the neighbours were weighted by the strength of functional correlations:

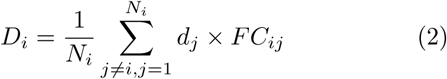

where *D*_*i*_ is collective deformation of functionally-defined neighbours of *i*-th node, *d* is node deformation, *N*_*i*_ is the degree of node *i* based on the group-level structural connectivity matrix, and *FC*_*ij*_ is the mean functional connectivity between nodes *i* and *j* across individuals. The summation excludes the deformation value of the node under consideration (*j* ≠ *i*). Note that we only included the nodes and their corresponding functional connections if they were structurally connected to the node under consideration (i.e., node *i*). Finally, the node deformation values were correlated with the collective deformation values of their neighbours weighted by functional connectivity

### Null models

Comparisons between deformation and connectivity were tested against two categories of null models. The first null model preserves spatial autocorrelation [1]. We first created a surface-based representation of the Cammoun atlas on the FreeSurfer *fsaverage* surface using the Connectome Mapper toolkit (https://github.com/LTS5/ cmp; [20]). We used the spherical projection of the *fsaverage* surface to define spatial coordinates for each parcel by selecting the vertex closest to the center-of-mass of each parcel. The resulting spatial coordinates were used to generate null models by applying randomly-sampled rotations and reassigning node values based on that of the closest resulting parcel (10,000 repetitions). The rotation was applied to one hemisphere and then mirrored for the other hemisphere. Importantly, this procedure was performed at the parcel resolution rather than the vertex resolution to avoid up-sampling the data and potentially altering the distribution of DBM values during the rotation and reassignment procedure.

The second null model preserves the spatial embedding (i.e., geometry) of the structural connectome [9, 31, 54]. Edges were first binned according to Euclidean distance. Within each length bin, pairs of edges were then selected at random and swapped [9]. The procedure was repeated 1,000 times, generating a population of rewired structural networks that preserve the degree sequence of the original network and approximately preserve the edge length distribution (i.e., spatial embedding) of the original network.

## RESULTS

### System-specific deformation

Schizophrenia-related deformation was first defined by contrasting the deformation maps of schizophrenia patients and healthy controls (Fig. S7 depicts the unthresholded deformation pattern). The statistically significant deformation pattern shown in Fig. 1A appears to mainly target brain regions associated with specific systems. To statistically assess if this is the case, we used a spatial permutation procedure. Voxels were first parcellated according to the multi-resolution Cammoun atlas [14]. Nodes were stratified according to their membership in Yeo resting-state networks [78] (Fig. 1B). We also investigated volume deformation relative to the cytoarchitectonic classification of human cortex according to the classic von Economo atlas [59, 69, 71, 72]. We used an extended version of the von Economo cytoarchitectonic partition with 7 classes [70] (Fig. 1C). Mean deformation values were first calculated within each resting-state network and cytoarchitectonic class for the finest parcellation resolution to ensure the best match to the partitions. To assess the extent to which these means are determined by the network partition, and not trivial differences in size, coverage or symmetry, we used a spherical projection null model that permutes network labels while preserving spatial autocorrelation [1] (referred to as “spin test” throughout the present report). Network labels were randomly rotated and mean deformation values were re-computed. The procedure was repeated 10,000 times and used to construct a distribution of network deformation means under the null hypothesis that regional deformation patterns are independent of affiliation with these large-scale systems.

**Figure 1.**
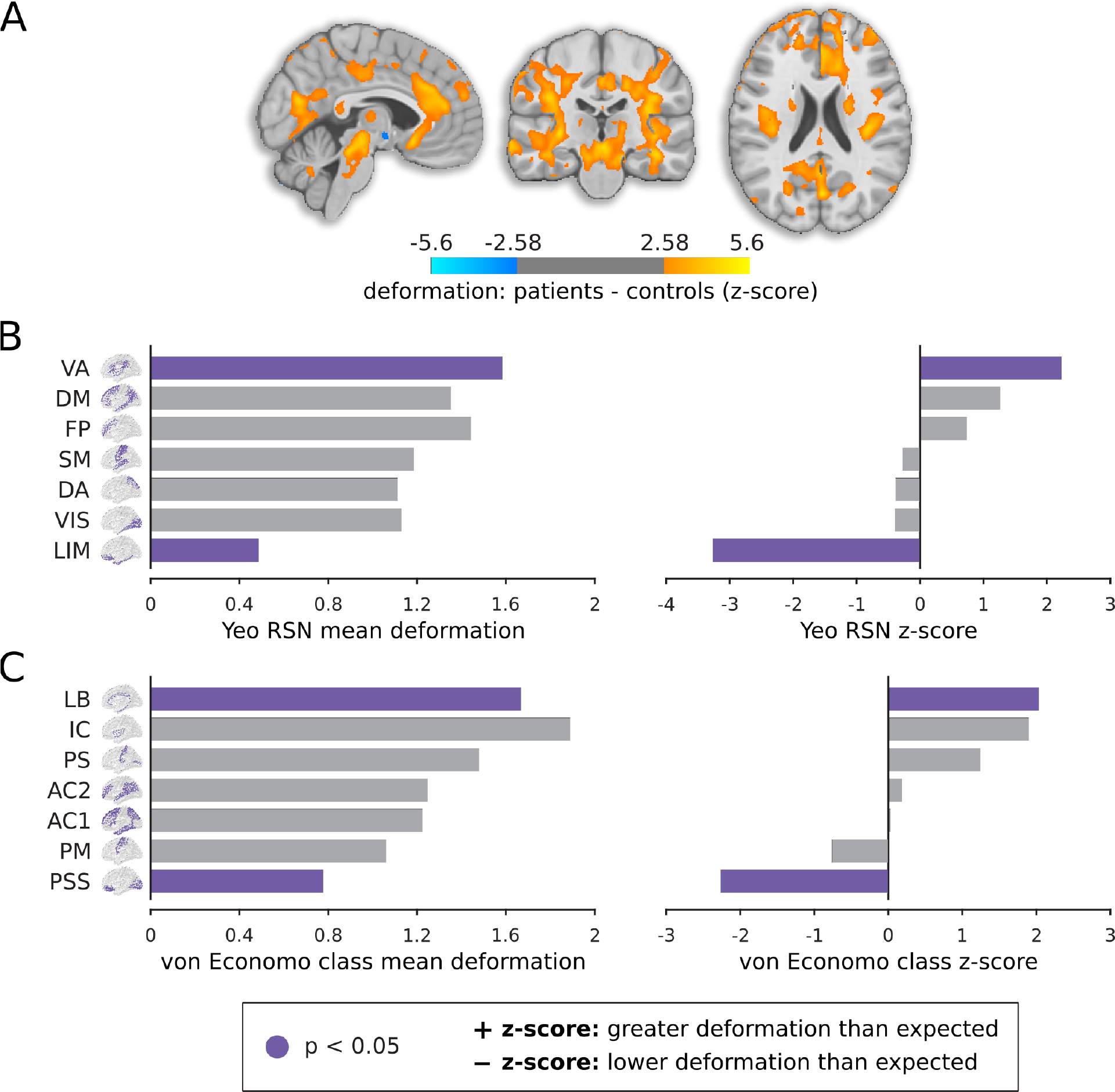
Deformation in schizophrenia patients compared to healthy controls. (A) Deformation-based morphometry maps of schizophrenia patients and controls were contrasted using two-sample *t*-tests. The *t*-statistics are converted to *z*-scores and displayed on an MNI template (MNI152_symm_2009a; (*x* = −4, *y* = −23, *z* = 22)). Greater *z*-scores correspond to greater deformation in schizophrenia patients relative to healthy controls. The maps are corrected for multiple comparisons by controlling the false discovery rate at 5% [7]. The deformation pattern is stratified into resting-state networks (RSNs) defined by Yeo and colleagues [78] (B) and into cytoarchitectonic classes of human cortex defined by the von Economo atlas [69–71] (C). The mean deformation score is calculated for each network. Network labels are then permuted, while preserving the spatial autocorrelation, and network-specific means are re-calculated. The procedure is repeated 10,000 times to generate a null distribution of network-specific deformation. The mean deformation for each RSN (B) and cytoarchitectonic class (C) and their corresponding *z*-scores relative to the null distribution generated by the spatial permuting procedure are depicted (10,000 spin tests; two-tailed). Positive *z*-score indicates greater deformation than expected, while negative *z*-score indicates lower deformation than expected. Yeo networks: DM = default mode, DA = dorsal attention, VIS = visual, SM = somatomotor, LIM = limbic, VA = ventral attention, FP = fronto-parietal. Von Economo classes: AC1 = association cortex, AC2 = association cortex, PM = primary motor cortex, PS = primary sensory cortex, PSS = primary/secondary sensory, IC = insular cortex, LB = limbic regions.

Fig. 1 shows the mean deformation values and the corresponding *z*-scores relative to the null distribution for each resting-state network (Fig. 1B) and cytoarchitectonic class (Fig. 1C). Consistent with the voxel-wise anatomical pattern, the ventral attention intrinsic network and the limbic cytoarchitectonic class display significantly greater deformation. Conversely, the limbic intrinsic network and primary/secondary sensory cytoarchitectonic class display significantly lower volume loss. Note the difference in the anatomical distributions of the so-called “limbic” systems between Yeo resting-state network and von Economo cytoarchitectonic class; the von Economo limbic class is focused on the cingulum, while the Yeo limbic network mainly includes orbitofrontal cortex and temporal poles [78]. Results were consistent across the five parcellation resolutions (Fig. S3).

### Structural and functional connectivity shape deformation

The fact that tissue volume loss is circumscribed by specific intrinsic networks suggests that deformation may be constrained by network architecture. We therefore directly test the possibility that the distribution of deformation patterns in schizophrenia is shaped by structural and functional connectivity. If this is the case, nodes strongly connected with high-deformation neighbours should display greater deformation, while nodes connected mainly with low-deformation neighbours should display less deformation (Fig. 2A).

**Figure 2.**
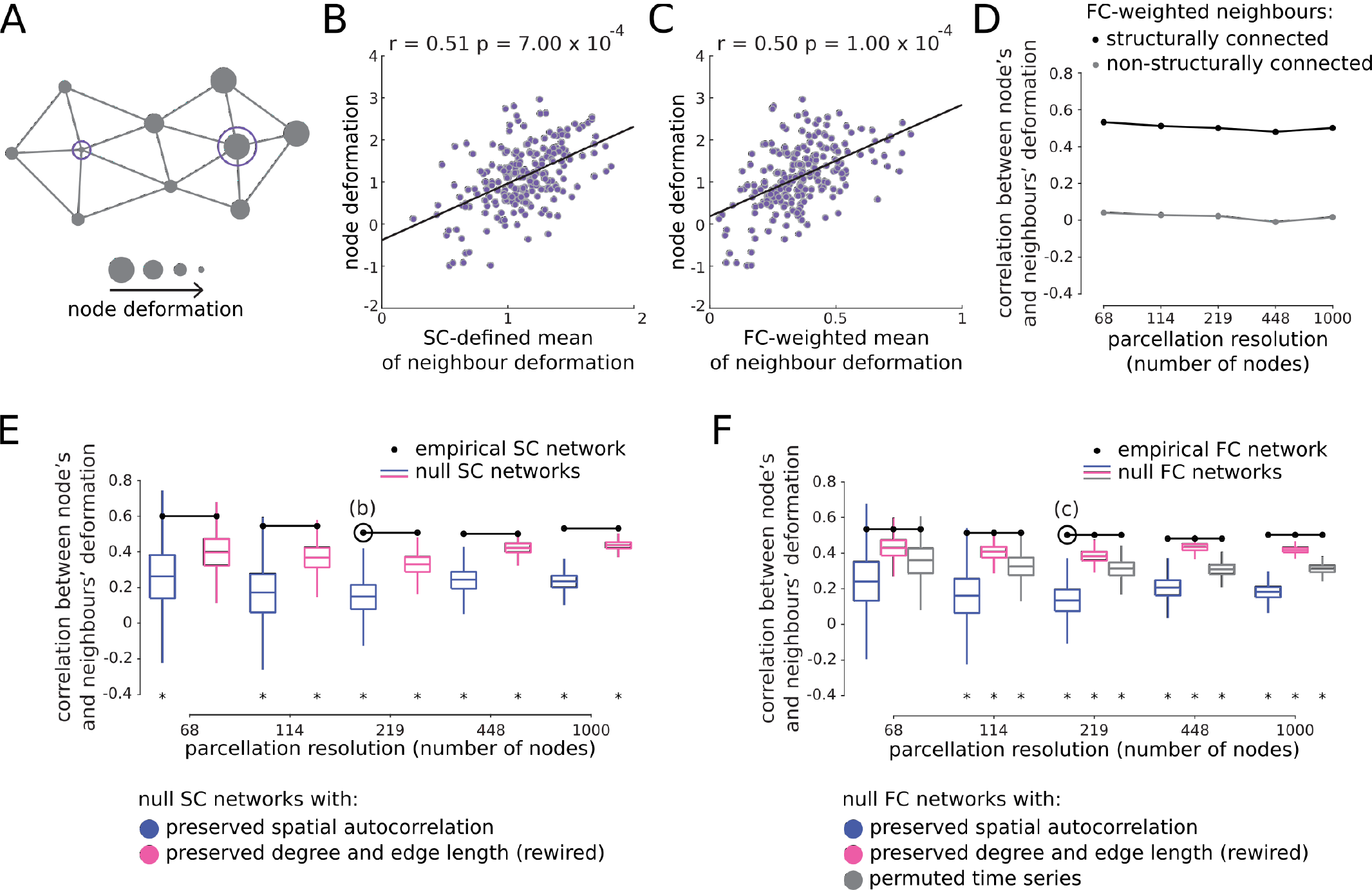
Network-dependent deformation. (A) Schematic of deformation of a node and its neighbours. If regional deformation depends on network connectivity, nodes connected to highly deformed neighbours will be more likely to be deformed themselves, while nodes connected to healthy neighbours will be less likely to be deformed. (B) The deformation of a node is correlated with the deformation of its neighbours defined by structural connectivity, estimated from an independently-collected diffusion-weighted MRI dataset. (C) The deformation of a node is correlated with the deformation of its neighbours weighted by functional connectivity, estimated from an independently-collected resting state functional MRI dataset. Both results are shown for the third resolution only (219 nodes). See Fig. S1 for results in all 5 resolutions, with and without the effect of distance regressed out from the data. (D) The correlations between node and neighbour deformation (weighed by functional connectivity) for structurally connected neighbours (black) were compared to the ones for non-structurally connected neighbours (grey). The associations remain only when considering structurally connected neighbours. (E,F) Two null structural models were constructed: (1) a spatial autocorrelation null model that projects nodes to a sphere and randomly rotates the sphere [1] (i.e., spin tests; 10,000 repetitions; blue); (2) a geometric null model that randomly rewires pairs of edges in the structural network, preserving the degree sequence and edge length distribution [9, 31, 54] (1,000 repetitions; pink). A null functional model was also constructed by randomly reassigning resting-state functional MRI time series to each node (1,000 repetitions; grey). The correlations between nodes’ and neighbours’ deformation values are depicted across the five resolutions using empirical structural and functional networks (black; e and f respectively), as well as the corresponding correlations using the null structural and functional networks. Statistically significant empirical correlations are indicated using an asterisk (*P* < 0.05; two-tailed).

To test this hypothesis, we investigate whether the deformation of a given brain region is correlated with the deformation of its connected neighbours. We derive group-level structural and functional connectivity networks from an independent sample of *N* = 70 participants (see *Methods and Materials*). For each network node, we correlate the local schizophrenia-related deformation value with the normalized sum of its structural or functional neighbours’ deformation values. For both structural and functional connectivity, the deformation of a node is significantly correlated with the deformation of its connected neighbours (resolution 3; *r* = 0.51, *P* = 7.00 × 10^−4^ and *r* = 0.50*, P* = 1.00 × 10^−4^ respectively (10,000 spins, two-tailed); Fig. 2B,C). The results are consistent across all five resolutions, suggesting that the effect does not trivially depend on how the network is defined (Fig. S1). The results are also consistent when we simultaneously include both structurally- and functionally-defined neighbours’ deformation estimates as predictors in a multiple regression model (resolution 3; adjusted-*r* = 0.53*, P* = 1.00 × 10^−4^ (10,000 spin tests; two-tailed); Table S1).

Can functional connectivity on its own - without reference to the underlying structural patterns - be used to predict deformation among nodes? To address this question, we computed the correlations between node and neighbour deformation (weighted by functional connectivity) for structurally connected neighbours (Fig. 2D; black). We also computed associations between nodes and non-structurally connected neighbours (weighted by functional connectivity) (Fig. 2D; grey). Fig. 2D shows that the associations remain only when considering structurally connected neighbours. We next investigated whether functional connectivity alone could explain the spatial patterning of tissue volume loss. We altered Equation 2 to include all neighbours (both with and without a structural connection) and measured the mean deformation of the functionally-weighted neighbours of each node. Table S2 shows the correlation between node and neighbour deformation, when all nodes are considered and weighted by their functional connectivity to the target node (column 4). The correlations are markedly reduced and are no longer statistically significant. Additionally, we measured the correlation between structurally- and functionally-defined neighbour deformation values in both cases. The mean deformation of structural and functional neighbours are highly correlated when only the structurally connected nodes are taken into account (Table S2, column 3 versus 5), suggesting that structural connectivity is the primary determinant of neighbour deformation. Altogether, the results demonstrate that functional connectivity between nodes is associated with their mutual deformation, but only if there exists an underlying white-matter connection.

We next seek to assess the extent to which these associations depend on network topology. To address this question, we constructed two null models: (1) a geometric null model that randomly rewires pairs of edges in the structural network, preserving the degree sequence and edge length distribution [9, 31, 54]; (2) a spatial autocorrelation null model that projects nodes to a sphere and randomly rotates the sphere [1] (see *Methods and Materials*). Fig. 2E,F shows the correlations between nodes’ and neighbours’ deformation values across the five resolutions using empirical structural and functional networks (black), as well as the corresponding correlations estimated using the two null models (10,000 repetitions for the spin test and 1,000 repetitions for the rewired null model). In both instances, the overall correspondence between node and neighbour deformation is greater in the empirical networks than the null networks (*P* < 0.05; two-tailed), with the exception of lowest parcellation resolution (only significant against the spin tests for SC-defined neighbours). We included an additional null functional model by randomly reassigning rs-fMRI time series to each node and re-estimating functional connectivity networks (1,000 repetitions). The results are consistent with the ones from the other two null models (Fig. 2F).

Finally, we ask whether deformation in a given node can be predicted using nodes that are not directly connected, but two or more synapses removed. To test this hypothesis, for each node we calculated the mean deformation of all the connected and not-connected neighbours. The neighbours were weighted either by the binary path length or the binary communicability between them [24]. In both instances the correlations were low (resolution3: *r* = −0.06 and *r* = −0.15) and were not statistically significant according to 10,000 spin tests (*P* = 0.9 and *P* = 0.1; two-tailed). Thus, we identify a strong influence of directly-connected neighbours, but find little evidence that neighbours further than one hop away make a reliable contribution to local deformation.

### Identifying the disease epicenter

Given that schizophrenia-related deformation depends on structural and functional connectivity, we next ask which brain regions might be the putative epicenters of the disease, analogous to the source of an epidemic. We hypothesized that a high-deformation node could potentially be an epicenter if its neighbours also experience high deformation (Fig. 3A). Nodes were ranked based on their deformation values and their neighbours’ deformation values in ascending order in two separate lists; we then identified nodes that were highly ranked in both lists (Fig. 3B) and assessed the significance of rankings using the spatial permutation testing (spin tests; 10,000 repetitions). The results depicted in Fig. 3C,D demonstrate that areas with significantly high mean rankings across both lists are primarily located in the bilateral cingulate cortices, consistent with previous literature [19]. To ensure that the effect is not driven by the number of connections (i.e., biased towards high-degree nodes), we normalized the deformation values of neighbours’ of a given node by its degree. The results were consistent across the 5 resolutions, indicating that the potential epicenter can be identified independent from the parcellation resolution (Fig. S2).

**Figure 3.**
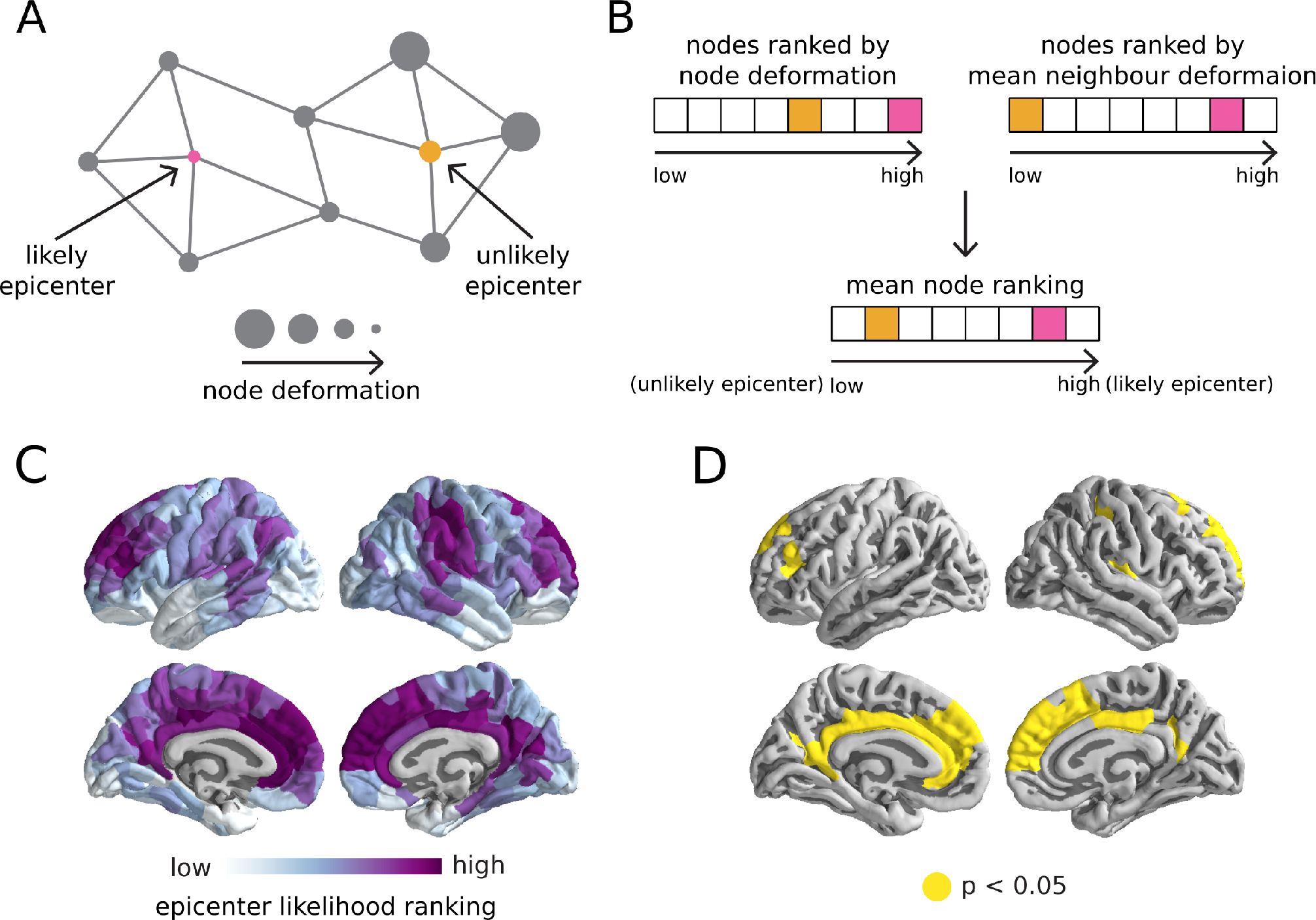
Epicenter identification. (A) We hypothesized that a high-deformation node could potentially be an epicenter if its neighbours also experience high deformation. (B) Nodes were ranked based on their deformation values and their neighbours’ deformation values in ascending order. Areas with high mean rankings across both lists are more likely to be an epicenter. (C) Mean rankings are depicted on the brain surface for the third parcellation resolution (See Fig. S2 for results in all 5 resolutions). (D) The statistical significance of the rankings was assessed using a spatial permutation testing approach (spin tests; 10,000 repetitions). Areas with significantly high rankings across both lists are primarily located in the bilateral cingulate cortices.

### Control analyses: spatial proximity, sex and medication, and replication

It is possible that local deformation correlates with the deformation of connected neighbours because spatially proximal nodes trivially exhibit greater co-deformation and greater connectivity. To rule out this possibility, we measured the mean Euclidean distance between a given node and its connected neighbours (centroid-to-centroid). We then regressed out Euclidean distance from both node deformation and its neighbours’ mean deformation and correlated the residual deformation values. As before, the deformation of a node was correlated with the deformation of its neighbours across resolutions (Fig. S1, bottom).

To further investigate the effect of spatial proximity, we excluded the spatially adjacent neighbours of the node under consideration from the calculation of mean neighbour deformation. More specifically, for a given node *i*, we first identify all the nodes that have any voxels spatially adjacent to node *i* (i.e., abutting node *i*) in the image used to parcellate the data. We then exclude all the spatially adjacent neighbours from Equations 1 and 2 and re-estimate the mean neighbour deformation for each node. We then assess the statistical significance of the correlation between node deformation and the mean deformation of non-adjacent neighbours using the null models described above. Consistent with the original analysis, the node and neighbour deformation values are significantly correlated even if only the non-adjacent neighbours are considered in the analysis (Fig. S4).

We next sought to investigate the effects of additional confounding factors, such as sex and antipsychotic medication dose [68]. We removed all participants for whom current antipsychotic medication dose was not available. For the remaining 87 participants (64 second-generation, 10 first-generation, 7 both, 6 non-medicated), we computed the chlorpromazine equivalent dosage [49, 76] and repeated the above analysis, controlling for medication dose in addition to age. We also repeated the analyses, controlling for sex in addition to age, for all the 133 participants. We find that the age- and sex-corrected and age- and medication-corrected deformation maps are highly correlated with the age-corrected maps (resolution 3; *r* = 0.95, *P* = 1.00 × 10^−4^ and *r* = 0.85, *P* = 1.00 × 10^−4^, respectively (10,000 spin tests; two-tailed); Fig. S5A). Moreover, we find that the relationship between neighbour connectivity and deformation is still present and remains independent of spatial proximity and choice of parcellation (Fig. S5B,C).

Finally, to ensure that the findings are not specific to the selected sample, we replicated the analyses in an independently-collected dataset (see *Methods and Materials*). Deformation patterns are significantly correlated between the discovery and replication datasets (Fig. S6A,B). Node deformation is positively correlated with its connected neighbours’ deformation and this relationship is significantly greater than in the null models (Fig. S6C). In addition, the putative epicenters of the disease are localized in the bilateral cingulate cortices, consistent with the discovery dataset (Fig. S6D).

## DISCUSSION

### Network patterning of deformation

The principal finding in the present investigation is that the deformation of an area is associated with the deformation of areas it is connected with. Interestingly, several recent studies found complementary results. Wannan and colleagues found that cortical thinning in schizophrenia was more likely among areas with greater anatomical covariance [73]. Di Biase and colleagues found that cortical thinning and reduction in local white matter anisotropy are anticorrelated in patients with schizophrenia [23]. Finally, Palaniyappan and colleagues found that patterns of cortical thinning are highly structured and systematically deviate from healthy anatomical covariance patterns [46]. Altogether, these results show that deformation patterns reflect network architecture, raising the possibility that connections drive pathology.

The structurally-guided deformation appears to accumulate in specific systems: the functionally-defined ventral attention network, and the cytoarchitectonically-defined limbic class. Moreover, given the structural connectome and the observed deformation pattern, we find that the likeliest epicenters are located in the cingulate cortices. These loci—situated at the intersection of the ventral attention/salience and default networks—are implicated in schizophrenia, including changes in tissue volume [25], structural connectivity [62, 79], activation [47, 48], and functional connectivity [12, 61, 75]. Interestingly, the same areas are frequently found to be affected in a wide range of mental illnesses, including schizophrenia, bipolar disorder, depression, addiction, obsessive-compulsive disorder and anxiety [32]. These regions may be particularly vulnerable because they are rich in von Economo neurons (VENs), which have also been associated with multiple psychiatric disorders including schizophrenia [2, 13, 16].

Despite the strong influence of directly-connected neighbours, we find little evidence that neighbours further than one hop away make a reliable contribution to local deformation. In other words, network connectivity plays an important role in shaping deformation but the influence is expressed mainly via direct connections and is more difficult to detect at longer path lengths. This result suggests the presence of additional, perhaps local, factors that shape deformation beyond the global connection patterns.

### A spreading hypothesis of schizophrenia?

Our results are broadly reminiscent of an emerging literature on the signature of network structure in neurodegeneration [26, 51]. In neurodegenerative diseases, transneuronal transport of toxic misfolded proteins is associated with cell death and atrophy [40, 43, 74]. As a result, patterns of atrophy in neurodegenerative diseases often resemble structural and functional network patterns [17], a result that has been reported in Alzheimer’s disease [35, 53, 60], Parkinson’s disease [77, 80, 81] and amyotrophic lateral sclerosis [58].

Do the present results support an analogous “spreading” hypothesis of schizophrenia? We have demonstrated that connectivity and deformation (i.e., reductions in brain tissue volume, but not necessarily cell death) are related in schizophrenia. Nevertheless, the evidence is circumstantial and it is impossible to disentangle the two. An alternative possibility is that systematic white matter disconnectivity - which we do not measure here - impedes normal trans-neuronal transport of trophic factors, leading to deformation among connected populations.

### Methodological considerations

We confirmed that the present results do not trivially depend on confounding factors such as parcellation resolution and spatial distance, but there are several technological factors that need to be taken into account as well. Structural connectivity, as estimated by diffusion weighted imaging, is prone to systematic false positives and false negatives [41, 66]. Connectomes generated by streamline tractography are undirected, so it is impossible to test the causal effect of connectivity on deformation. Likewise, we cannot directly measure neuronal malformation *in vivo*; we instead operationalized deformation as changes in tissue volume density.

More generally, the current results are correlational: it is impossible to infer whether deformation drives structural and/or functional disconnectivity reported in previous literature or vice versa, or whether there is another underlying mechanism that independently potentiates both tissue loss and disconnectivity [21, 27, 30, 55, 56]. We opted to use a high-quality healthy young control dataset as a “gold standard” for structural and functional connectivity. Our rationale for using connectivity from healthy controls was that tissue volume loss in patients might compromise connectivity among affected regions. As a result, connection patterns derived from patients may misrepresent the architectural foundation for the distributed deformation patterns we observe in chronic patients. However, this approach ignores extensive changes in structural and functional connectivity that are well-documented in schizophrenia [21, 30]. For example, it is possible that deterioration in white matter connectivity obstructs or reroutes the spread of pathology. How changes in tissue volume and disconnectivity mutually evolve over the time course of the disease remains an exciting open question; application of the present methodology to longitudinal samples will help to resolve the temporal evolution of these effects.

Finally, there is limited information available on medication history or adherence of the patients, making it difficult to assess whether medication influences the results. Our control analyses on a subset of patients indicate that the network effects reported here are unlikely to be correlated with chlorpromazine equivalent antipsychotic medication dose. Other groups have recently reported similar relationships between anatomical covariance networks and changes in cortical thickness in first-episode psychosis, chronic schizophrenia and treatment-resistant schizophrenia [73], suggesting that the observed effect is likely to be present earlier in the disease and in drug-naive patients.

## ACKNOWLEDGMENTS and DISCLOSURES

This research was undertaken thanks in part to funding from the Canada First Research Excellence Fund, awarded to McGill University for the Healthy Brains for Healthy Lives initiative. BM acknowledges support from the Natural Sciences and Engineering Research Council of Canada (NSERC Discovery Grant RGPIN #017-04265) and from the Fonds de Recherche du Québec - Santé (FRSQ, Chercheur Boursier). GS acknowledges support from the Natural Sciences and Engineering Research Council of Canada (NSERC) and from the Healthy Brains for Healthy Lives (HBHL) initiative at McGill University. CM acknowledges support from the Canadian Institutes of Health Research (CIHR) and from the Healthy Brains for Healthy Lives (HBHL) initiative at McGill University. GAD is supported in part by funding provided by Brain Canada, in partnership with Health Canada, for the Canadian Open Neuroscience Platform initiative. PH acknowledges Swiss National Science Foundation grant #156874. ML acknowledges support from the Canadian Institutes of Health Research (CIHR, 68961). ML has received a Salary award from the Fonds de Recherche du Québec Santé (FRSQ) and a James McGill Professorship. ML reports grants from Otsuka Lundbeck Alliance, personal fees from Otsuka Canada, personal fees from Lundbeck Canada, grants and personal fees from Janssen, and personal fees from MedAvante-Prophase, outside the submitted work. The authors report no other biomedical financial interests or potential conflicts of interest.

The authors thank Dr. Alessandra Griffa for data collection and processing of healthy structural and functional networks and providing insightful comments on the manuscript. The authors also thank Dr. Andrew Zalesky for providing insightful comments and feedback on the manuscript and Dr. František Váša for sharing the von Economo cytoarchitectonic atlases.

Please note that the present report has been posted on the *bioRxiv* preprint server (https://www.biorxiv.org/).

## Supplemental Information

### Deformation-Based Morphometry (DBM)

Automated pre-processing was performed using the *minc-bpipe-library* pipeline (https://github.com/CobraLab/minc-bpipe-library) on each T1-weighted MRI scan. The outputs were visually inspected and quality-controlled. A group-average template for the remaining participants, both healthy controls and patients, was built using the ANTs multivariate template construction tool [4] (https://github.com/CobraLab/documentation/wiki/ ANTs-Multivariate-Template-Construction).

Local changes in brain tissue volume density were calculated using deformation-based morphometry (DBM; [3]). Regional DBM values are estimated from the deformation applied to each voxel to non-linearly register each MRI scan to a standard template and can be used as measures of tissue loss (termed as *deformation* in this manuscript) or tissue expansion [15, 18, 39, 64, 80]. In the present study, we used ANTs multivariate template construction pipeline [4] (*antsMultivariateTemplateConstruction2.sh*) to measure the DBM values. This pipeline produces a population average through the iterative estimation and application of affine non-linear warps to a starting rigid model (here the MNI ICBM symm 2009c model). The final iteration non-linear transformation of each structural brain image to the unbiased template image produced during the registration process is used as a deformation based map for each subject in the template space. A deformation map quantifies the amount of displacement of each voxel in each direction in the 3-dimensional template space that was required to transform each brain image from subject to template space. The local change in tissue density is then estimated as the derivative of the displacement of a given voxel in each direction by calculating the determinant of the Jacobian matrix of displacement [80]. Thus, no change in volume is given with 1 (i.e., no displacement relative to template), tissue expansion is given with a value between 0 and 1 (i.e., subject image was shrunk to be transformed to template space), and tissue loss or deformation is given with a value larger than 1 (i.e., subject image was expanded to be transformed to template space). For easier interpretation of the changes in volume density, we calculated the logarithm of the determinant, such that no change is given by 0, tissue loss (i.e., deformation) is given by a positive value, and tissue expansion is given by a negative value.

Following the deformation procedure, the maps were blurred at twice the resolution of the input images (i.e., 2 mm full-width/half-maximum) and the non-brain tissue was removed from each brain using FreeSurfer image analysis suite (release v6.0.0; http://surfer.nmr.mgh.harvard.edu/). The voxel-wise data were then extracted for each participant for further analysis.

DBM was chosen over voxel-based morphometry (VBM) because the latter requires more extensive spatial smoothing and therefore has lower spatial resolution. This is important for the present study because we sought to correlate deformation values in brain regions with deformation values in neighbouring regions. There is also some evidence to suggest that DBM is more sensitive to grey matter changes in subcortex [11, 57], which have been demonstrated in schizophrenia [67].

### Healthy structural and functional networks

Structural and functional connectivity data, collected from 70 healthy individuals at Lausanne University Hospital in Switzerland (age 28.8 ± 9.1 years, 27 females) on a 3T scanner and described in detail elsewhere [33, 45] (http://doi.org/10.5281/zenodo.2872624), were used to construct high-quality reference brain networks of a healthy population. In brief, deterministic streamline tractography was used to construct structural connectivity matrices for each healthy individual from their diffusion spectrum imaging (DSI) data at each of the five parcellation resolutions. Each pair-wise structural connection was weighted by fiber density. In order to correct for any size differences between the two brain areas as well as the inherent bias towards longer fibers in tractography process, the fiber density of structural connections was estimated as the number of streamlines normalized by the mean surface area of the two regions and the mean length of streamlines connecting them [34, 45]. Finally, a binary group-average structural connectivity matrix was generated using a consensus approach preserving the edge length distribution in individual participants [8, 10, 44, 45].

Furthermore, functional data were collected for the same 70 participants using eyes-open resting-state functional MRI (rs-fMRI) scans and were pre-processed as described in [45]. Briefly, fMRI volumes were first corrected for physiological variables, including regression of white matter, cerebrospinal fluid and motion (three translations and three rotations, estimated by rigid body co-registration). BOLD time series were then lowpass filtered (temporal Gaussian filter with full width half maximum equal to 1.92 s). The first four time points were excluded from subsequent analysis. High motion frame censoring (“scrubbing”) was performed as described by Power and colleagues [52]. Functional connectivity between pairs of brain regions was estimated as zero-lag Pearson correlation coefficient between rs-fMRI time series of the two regions for each individual. The group-average functional connectivity matrix was estimated as the mean connectivity of pair-wise connections across individuals. We retained negative functional connections and used unthresholded functional connectivity data for the original analyses reported in the main text. However, we repeated the analyses after thresholding the negative connections (i.e., assigning them to zero) to rule out the possibility of any confounding effects. The results remained the same when we removed the negative functional connections both at the individual subject level and at the group-average level (Fig. S8).

---

**TABLE S1.** Multiple regression model. The relationship between deformation of a node and its neighbours for schizophrenia patients was estimated using a multiple regression model, where deformation values of structurally- and functionally-defined neighbours were simultaneously included as predictors. *P*-values are estimated from 10,000 rotational permutations (i.e., spin tests; two-tailed).

|  | **n=68** | **n=114** | **n=219** | **n=448** | **n=1000** |
| --- | --- | --- | --- | --- | --- |
| inter-node distance not regressed:<br>adjusted-$r$<br>$P$-value | 0.59<br>$1.61 \times 10^{-2}$ | 0.56<br>$1.00 \times 10^{-3}$ | 0.53<br>$1.00 \times 10^{-4}$ | 0.52<br>$1.00 \times 10^{-4}$ | 0.55<br>$9.99 \times 10^{-5}$ |
| inter-node distance regressed:<br>adjusted-$r$<br>$P$-value | 0.56<br>$1.89 \times 10^{-2}$ | 0.51<br>$3.70 \times 10^{-3}$ | 0.43<br>$2.00 \times 10^{-4}$ | 0.44<br>$2.00 \times 10^{-4}$ | 0.51<br>$9.99 \times 10^{-5}$ |

**TABLE S2.** Influence of functional connectivity on the spatial patterning of tissue volume loss. Correlations between node’s and FC-weighted mean neighbours’ deformation values for the structurally connected neighbours are provided in column 2 (across five parcellation resolutions given in column 1). These correlations are markedly reduced when the mean neighbour deformation is defined by all FC-weighted neighbours, irrespective of structural connectivity (column 4). Additionally, the correlations between structurally- and functionally-defined neighbour deformation values were estimated in both cases. The mean deformation of structural and functional neighbours are highly correlated when only the structurally connected nodes are taken into account (column 3 versus 5).

| **Resolution** | **Structurally connected neighbours** |  | **All neighbours** |  |
| --- | --- | --- | --- | --- |
|  | Correlation between node's and neighbours' deformations (FC weight) | Correlation between SC neighbours' (no weight) and SC neighbours' deformations (FC weight) | Correlation between node's and all neighbours' deformations (FC weight) | Correlation between SC neighbours' (no weight) and all neighbours' deformations (FC weight) |
| n = 68 | 0.53 | 0.80 | 0.32 | 0.16 |
| n = 114 | 0.51 | 0.72 | 0.26 | -0.005 |
| n = 219 | 0.50 | 0.76 | 0.17 | 0.02 |
| n = 448 | 0.48 | 0.74 | 0.07 | 0.02 |
| n = 1000 | 0.50 | 0.74 | 0.06 | 0.07 |

**Figure S1.**
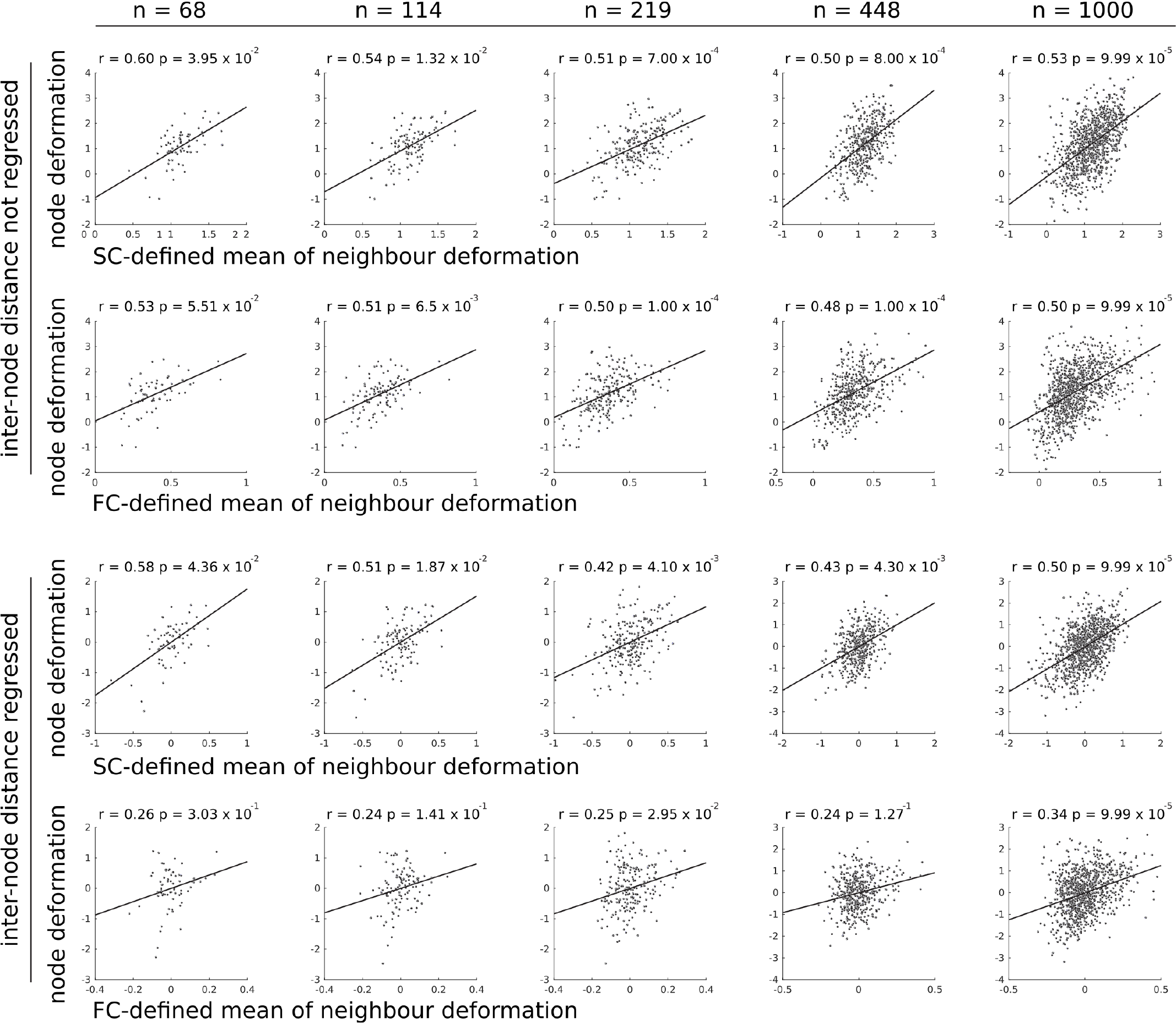
Network-dependent deformation (contrasting schizophrenia patients and controls) Deformation is defined using a general linear model to identify regions where deformation is different between patients with schizophrenia and controls (controlling for age). The deformation of a node is correlated with the deformation of connected neighbours, defined by structural connectivity (SC) and functional connectivity (FC). Top and bottom: Results are shown with and without removing the effect of spatial proximity. Left to right: Correlations are shown for five progressively finer anatomical parcellations [14]. *P*-values are obtained from 10,000 spatial permutation tests (i.e., spin tests; two-tailed).

**Figure S2.**
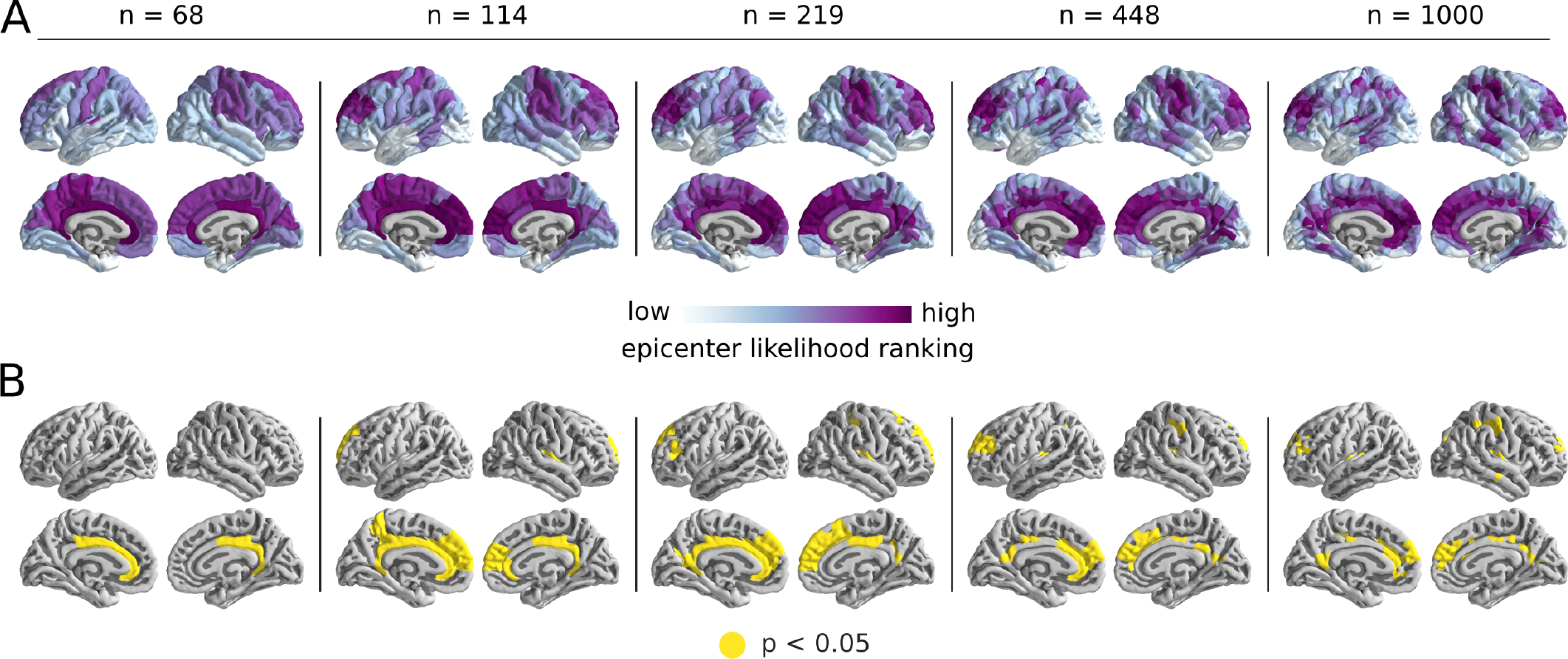
Epicenter analysis across parcellation resolutions. (A) The epicenter identification analysis was repeated across five progressively finer anatomical parcellations. The nodes were ranked based on their deformation values and their neighbours’ deformation values in ascending order. The mean rankings are depicted on the brain surface. (B) The areas with significant, mean highest rankings are consistently located in the bilateral cingulate cortices across the parcellations (10,000 spin tests).

**Figure S3.**
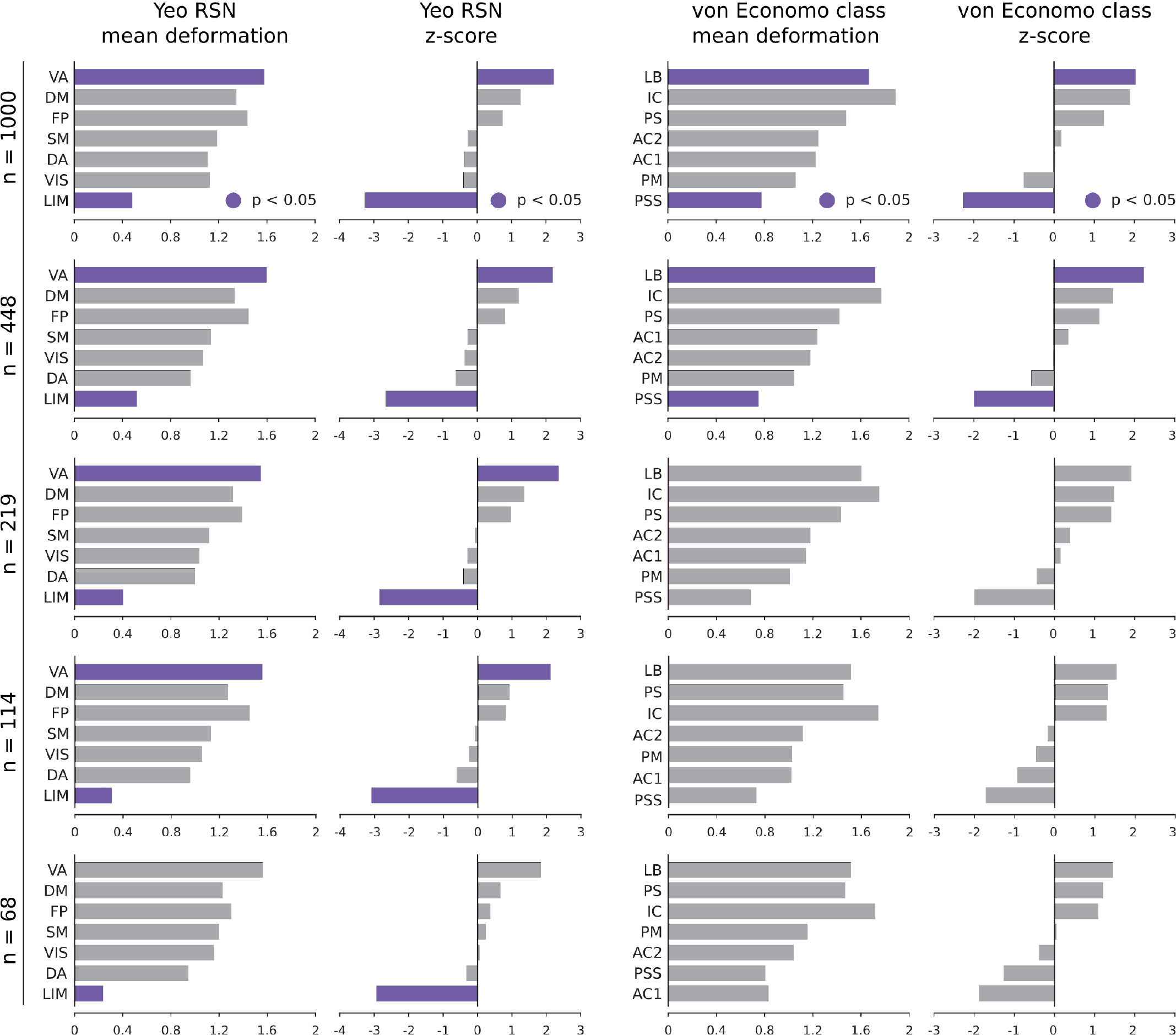
System-specific deformation. The mean deformation for Yeo resting state networks [78] and von Economo cytoarchitectonic classes [69–71] and their corresponding z-scores relative to the null distribution generated by the 10,000 spatial permutations (i.e., spin tests; two-tailed) are depicted across five progressively finer anatomical parcellations. Yeo networks: DM = default mode, DA = dorsal attention, VIS = visual, SM = somatomotor, LIM = limbic, VA = ventral attention, FP = fronto-parietal. Von Economo classes: AC1 = association cortex, AC2 = association cortex, PM = primary motor cortex, PS = primary sensory cortex, PSS = primary/secondary sensory, IC = insular cortex, LB = limbic regions.

**Figure S4.**
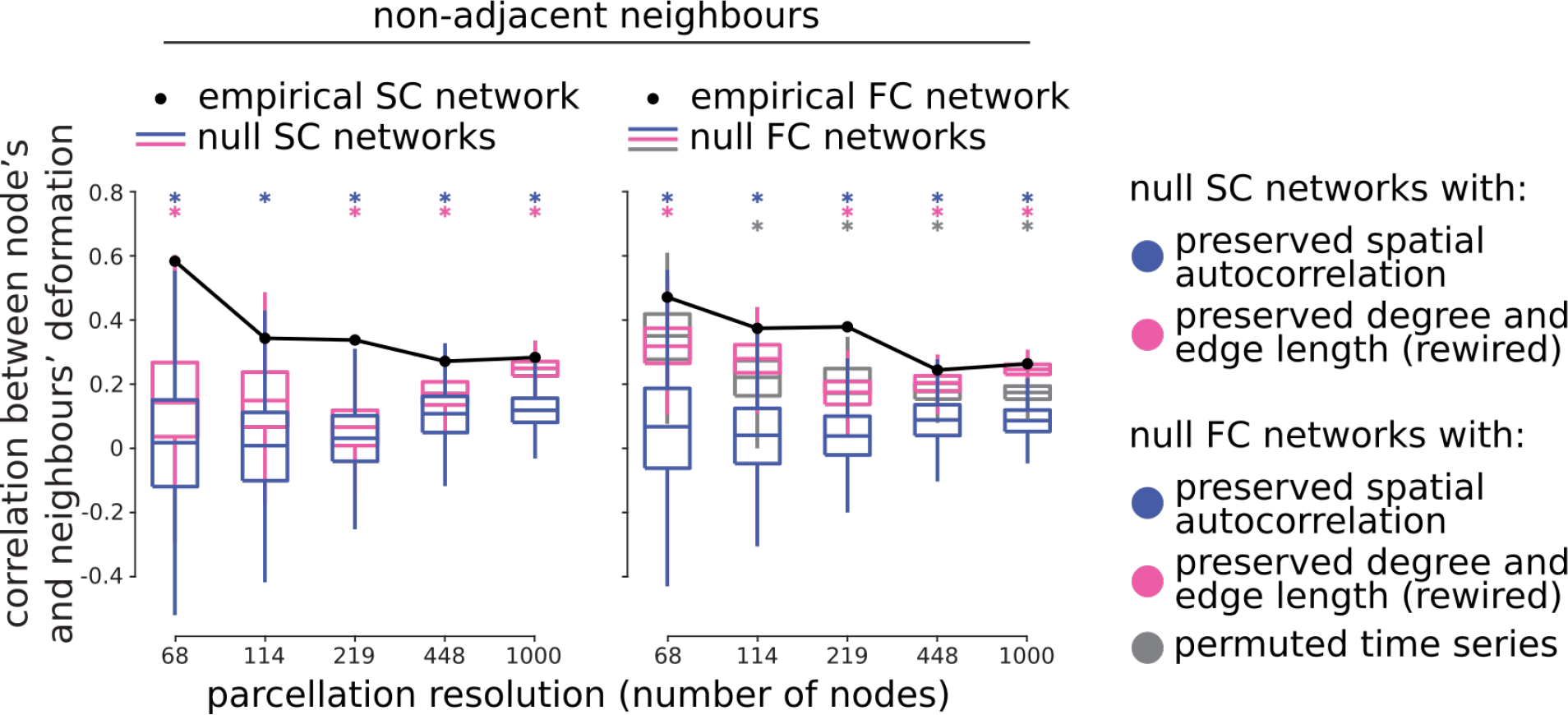
The relationship between a node’s and its spatially non-adjacent neighbours’ deformation values. Node deformation was correlated with the mean deformation of spatially non-adjacent neighbours (i.e., non-abutting neighbours) across five parcellation resolutions, such that the spatially adjacent neighbours of a given node were excluded from the calculation of its mean neighbour deformation. The statistical significance of the correlations was assessed against the structural and functional null models: (1) a spatial autocorrelation null model that projects nodes to a sphere and randomly rotates the sphere [1] (i.e., spin tests; 10,000 repetitions; blue); (2) a geometric null model that randomly rewires pairs of edges in the structural network, preserving the degree sequence and edge length distribution [9, 31, 54] (1,000 repetitions; pink); (3) A null functional model was also constructed by randomly reassigning resting-state functional MRI time series to each node (1,000 repetitions; grey). Asterisks, colored according to the corresponding null model, indicate significant empirical correlations (*P* < 0.05; two-tailed).

**Figure S5.**
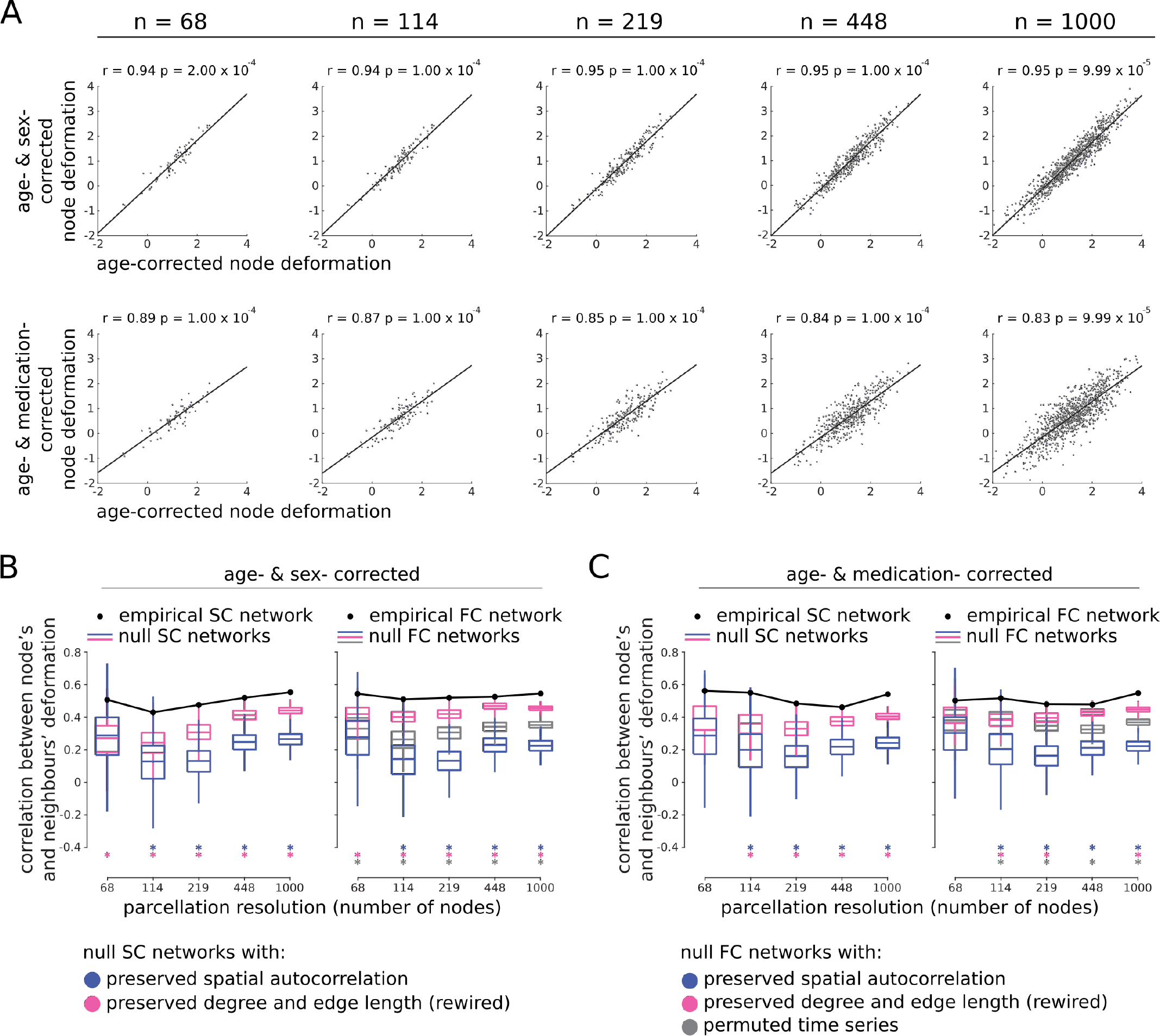
Control analysis: effects of confounding factors. (A) Deformation is defined using a general linear model controlling for age and sex (top) and for age and medication (bottom; for a subset of patients (*N* = 87) for whom the chlorpromazine equivalent antipsychotic medication dose information was available). The age- and sex-corrected and age- and medication-corrected deformations are highly correlated with age-corrected deformation pattern. (B) The correlations between node’s and neighbours’ age- and sex-corrected deformation values are depicted across the five parcellation resolutions using empirical structural and functional networks (black). The statistical significance of the correlations was assessed against the structural and functional null models: (1) a spatial autocorrelation null model that projects nodes to a sphere and randomly rotates the sphere [1] (i.e., spin tests; 10,000 repetitions; blue); (2) a geometric null model that randomly rewires pairs of edges in the structural network, preserving the degree sequence and edge length distribution [9, 31, 54] (1,000 repetitions; pink); (3) A null functional model was also constructed by randomly reassigning resting-state functional MRI time series to each node (1,000 repetitions; grey). Asterisks, colored according to the corresponding null model, indicate significant empirical correlations (*P* < 0.05; two-tailed). (C) The correlations between node’s and neighbours’ age- and medication-corrected deformation values are depicted across the five resolutions using empirical structural and functional networks (black), as well as the correlations obtained from the null structural and functional networks. The significance of the empirical correlations is shown using asterisks colored according to the corresponding null model (*P* < 0.05; two-tailed).

**Figure S6.**
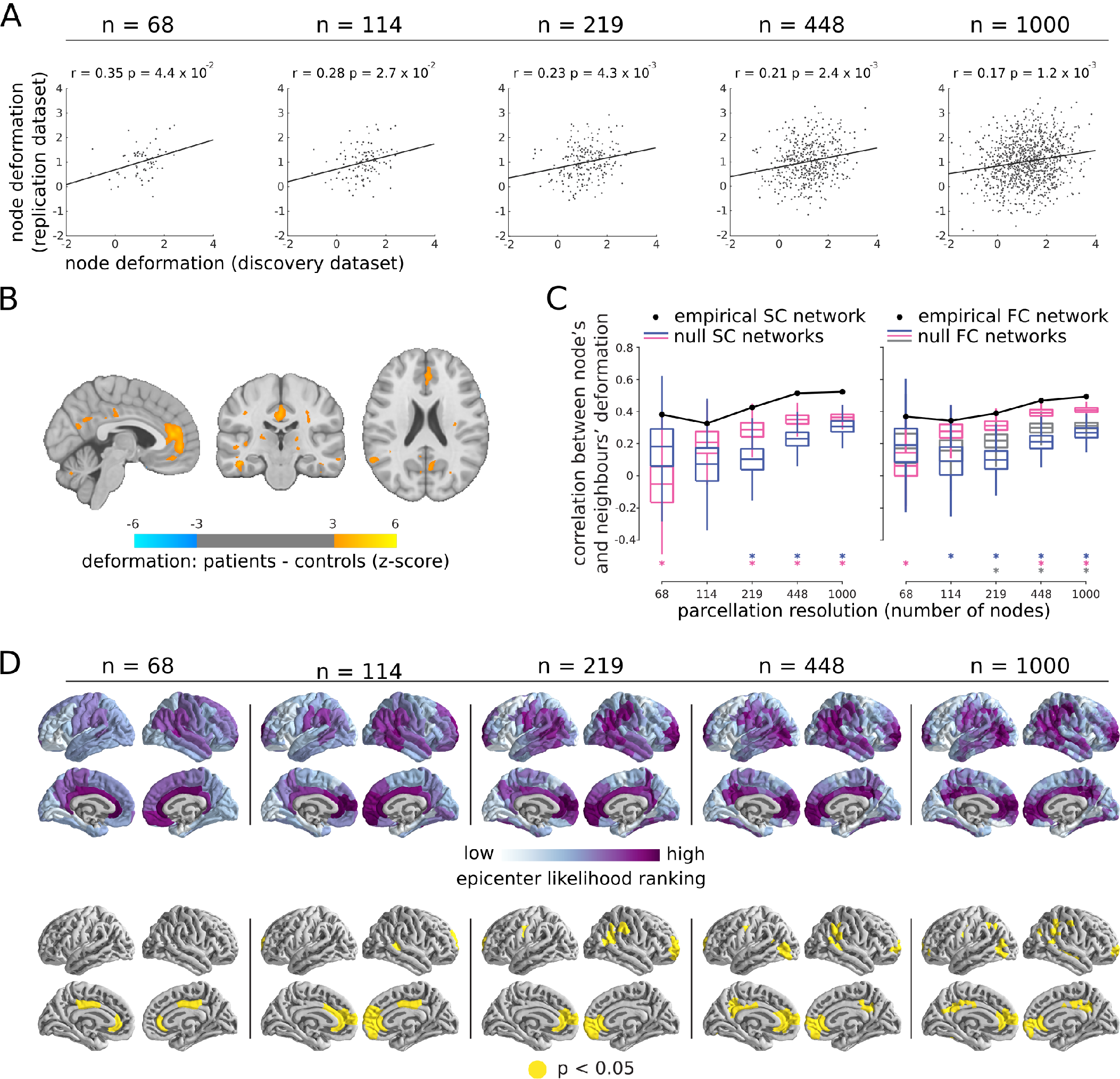
Replication dataset. (A) Schizophrenia-related deformation pattern was estimated using an independent dataset with *N* = 108 individuals with schizophrenia and *N* = 68 healthy controls. The deformation pattern in replication dataset is significantly correlated with the deformation pattern obtained from the discovery dataset across five progressively finer parcellation resolution. (10,000 spin tests; two-tailed). (B) The deformation pattern is depicted for the replication dataset. The *t*-statistics are converted to *z*-scores and displayed on an MNI template (MNI152_symm_2009a; (*x* = −4, *y* = −23, *z* = 22)). Greater *z*-scores correspond to greater deformation in schizophrenia patients relative to healthy controls. The maps are corrected for multiple comparisons by controlling the false discovery rate at 5% [7]. (C) The correlations between node’s and neighbours’ deformation values are depicted across the parcellations using empirical structural and functional networks (black). The statistical significance of the correlations was assessed against the structural and functional null models: (1) a spatial autocorrelation null model that projects nodes to a sphere and randomly rotates the sphere [1] (i.e., spin tests; 10,000 repetitions; blue); (2) a geometric null model that randomly rewires pairs of edges in the structural network, preserving the degree sequence and edge length distribution [9, 31, 54] (1,000 repetitions; pink); (3) A null functional model was also constructed by randomly reassigning resting-state functional MRI time series to each node (1,000 repetitions; grey). Asterisks colored according to the corresponding null model, indicate significant empirical correlations (*P* < 0.05; two-tailed). (D) The epicenter identification analysis was applied to the replication dataset across the parcellations. The nodes were ranked based on their deformation values and their neighbours’ deformation values in ascending orders. The mean rankings are depicted on the brain surface. The areas with significant, mean highest rankings are consistently located in the bilateral cingulate cortices across the parcellations (10,000 spin tests).

**Figure S7.**
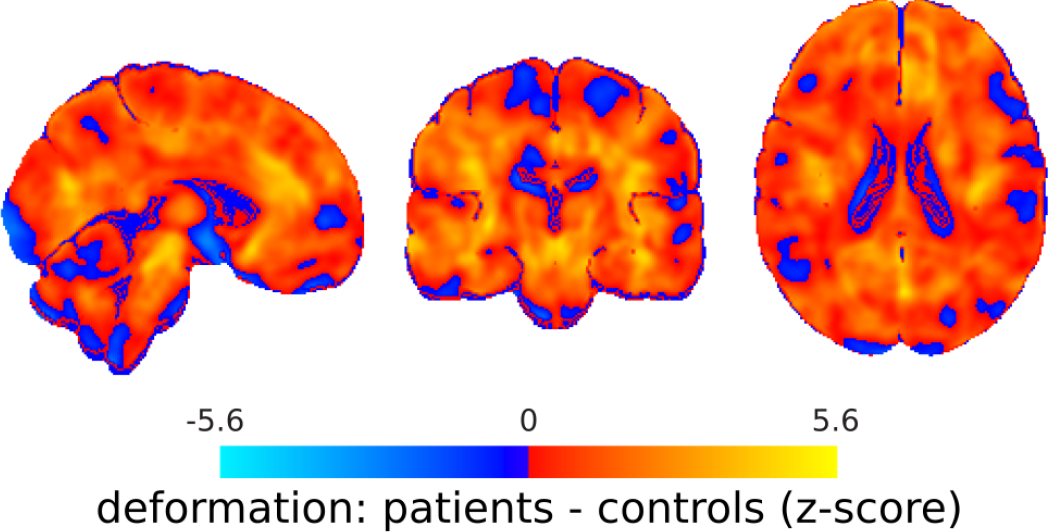
Unthresholded deformation pattern. Unthresholded schizophrenia-related deformation is depicted for the NUSDAST dataset. The deformation pattern was identified by contrasting the deformation maps of schizophrenia patients and healthy controls, using a mass-univariate analysis (i.e., two-tailed two-sample t-test; controlling for age) of voxel-wise differences between DBM values of the two groups.

**Figure S8.**
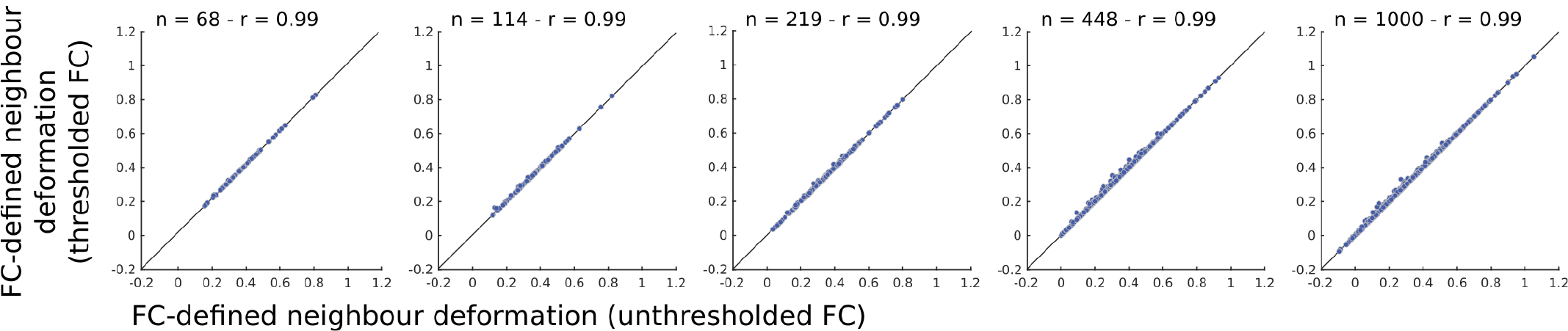
Effect of thresholded functional connectivity on neighbour deformation. Group-average functional connectivity network was used to estimate mean neighbour deformation (Equation 2). Negative functional connections were retained and unthresholded functional connectivity data was used for the original analyses reported in the manuscript. However, the analysis was repeated after thresholding the negative connections (i.e., assigning them to zero) to rule out the possibility of any confounding effects. Mean neighbour deformation pattern calculated using the thresholded functional connections is highly correlated with the mean neighbour deformation estimated from the unthresholded functional connectivity data across the parcellation resolutions. Functional connectivity data was thresholded at the group-average level in the results depicted here (the results are consistent when the negative functional connections are removed at the individual subject level).

